# Evolution of hominin detoxification: Neanderthal and modern human AHR respond similarly to TCDD

**DOI:** 10.1101/445239

**Authors:** Jac M.M.J.G. Aarts, Gerrit M. Alink, Henk J. Franssen, Wil Roebroeks

## Abstract

In studies of hominin adaptations to fire use, the role of the aromatic hydrocarbon receptor (AHR) in the evolution of hominin detoxification, especially regarding toxic smoke components, has been highlighted, including statements that the modern human AHR is significantly better at dealing with smoke toxins. We compared the AHR-controlled induction of cytochrome P450 1A1 (CYP1A1) in cultured cells transfected with an Altai-Neanderthal respectively a modern human reference AHR expression construct, and exposed to 2,3,7,8-tetrachlorodibenzo-*p*-dioxin (TCDD). We compared the complete AHR mRNA sequences including the untranslated regions (UTRs), maintaining the original codon usage, in HeLa human cervix epithelial adenocarcinoma cells. Our experiments complement a previous study of the AHR coding region optimized for mammalian codon usage and expressed in rat cells. Our results show no significant difference in CYP1A1 induction by TCDD between Neanderthal and modern human AHR (instead of a previously reported 150-1000 times difference range before), demonstrating that expression in a homologous cellular background is of major importance. The two dose-response curves almost coincide, except for a higher extrapolated maximum induction level for the Neanderthal AHR, possibly caused by a 5’-UTR G-variant known from modern humans (rs7796976). Our results are strongly at odds with a major role of the modern human AHR in the evolution of hominin detoxification of smoke components and consistent with our previous study which concluded that efficient detoxification alleles are more dominant in ancient hominins than in modern humans based on 18 relevant genes in addition to AHR.

## INTRODUCTION

The use of fire is a defining characteristic of the human lineage, with pyrotechnology one of the most powerful tools developed during human evolution. Fire afforded humans with (well-known) benefits affecting many domains of the human niche, including food preparation and diet, defense against predators, thermoregulation and social interaction (Carmody *et al.*, 2009; Wiessner, 2014; Wrangham, 2009).

However, pyrotechnology also came with (lesser known) costs (Henry *et al.*, 2018), including toxicological ones (Aarts *et al.*, 2016). The utilization of fire fueled with wood or other types of biomass on a daily basis by prehistoric hominins implied frequent exposure to toxic components of smoke, to which polycyclic aromatic hydrocarbons (PAHs) contribute importantly (Freeman *et al.*, 1990). In view of the well-known reproduction-toxic effects of tobacco smoke (Dechanet *et al.*, 2011; DeMarini, 2004) and the very similar composition of smoke from burning any type of biomass (Mishra *et al.*, 2005; Naeher *et al.*, 2007), the capacity to detoxify smoke toxicants is a crucial fitness factor inferred to have been under positive genetic selection in hominin populations ever since they started using fire regularly.

The chronology of fire use and fire production however is unclear: some scholars advocate a long chronology, with routine fire use starting with *Homo erectus*, more than two million years ago (Wrangham, 2009; Wrangham *et al.*, 2010). Based on archaeological data, others see consistent fire use as a significantly later phenomenon, from around 350,000 years ago onward (Roebroeks *et al.*, 2011; Shimelmitz *et al.*, 2014), while regular fire production has even been inferred to be a modern human accomplishment only (Sandgathe *et al.*, 2011) (but see (Sorensen, 2017; Sorensen *et al.*, 2018)). In an attempt to shed independent light on the debated chronology, Aarts *et al.* (2016) hypothesized that frequent exposure to toxic compounds occurring in smoke would have resulted in genetic adaptations in genes involved in detoxification of these toxicants. They analyzed 36 genetic variants in a comprehensive set of 19 relevant genes in Neanderthal, Denisovan, and (pre)historic and extant anatomically modern human genomes. This showed that archaic hominins predominantly possessed protective gene variants, as do extant chimpanzees. In contrast, the number of less protective, “high-risk” alleles has increased in modern humans. This pattern starts with the earliest modern human genome, of the 45 kya Ust’-Ishim individual (Fu *et al.*, 2014), and is suggestive of a deterioration in the ability to deal with smoke toxicants in modern humans, starting long before the emergence of agriculture. The more efficient detoxification of smoke toxins by Neanderthals and Denisovans was apparently hitchhiking on old primate mechanisms likely involved in dealing with plant toxins, and possibly in balancing certain photo-oxidation products induced by exposure to UV light.

Cytochrome P450 1A1 (CYP1A1) plays an important, dualistic role in the defense against PAH intoxication, being involved in both the generation of mutagenic radical-type reaction intermediates, and in phase 1 detoxification of PAHs (Divi *et al.*, 2014; Nebert *et al.*, 2013). The CYP1A1 gene is under transcriptional control of the aryl hydrocarbon receptor (AHR) (Corchero *et al.*, 2001; Nukaya *et al.*, 2009), a ligand-activated key regulator of many detoxification genes (Köhle *et al.*, 2007). In addition to dioxins and dioxin-like compounds such as 2,3,7,8-tetrachlorodibenzo-*p*-dioxin (TCDD), and PAHs such as benzo[*a*]pyrene (BaP), the AHR is activated by a very diverse range of agonist molecules (Denison *et al.*, 2011). Therefore, the AHR gene variant and the resulting detoxification phenotype of ancient hominins is an important aspect of the evolution of smoke detoxification.

Starting from an evolutionary hypothesis comparable to Aarts *et al.* (2016), Hubbard *et al.* (2016) focused on the aryl hydrocarbon receptor (AHR) gene, for which humans display a fixed difference (Val381) from the Neanderthal and Denisovan (Ala381) ancestral variant. Hubbard *et al.* (2016) performed an analysis of the modern human versus the Neanderthal AHR in rat cells and concluded that the Neanderthal AHR induced CYP1A1 mRNA expression with an EC_50_ between 150- and 1000-fold lower than observed with the human AHR. While acknowledging that the potential fitness effects associated with fixation of the V381 variant in humans, if any, are not known, Hubbard *et al.* (2016, page 2654) emphasized that “one plausible explanation is that this mutation provides a degree of protection against the deleterious effects of toxic environmental AHR ligand exposure”. In their view, having this mutation made a dramatic, hundred-fold to as much as a thousand-fold difference (Caspermeyer, 2016: page 2767) which “may have represented a gene-culture evolutionary advantage for humans” (Hubbard *et al.* 2016: page 2655).

Here we follow their approach and focus on the AHR, but with some important differences in experimental set-up based on the literature on AHR biology and function (Ema *et al.*, 1994; Moriguchi *et al.*, 2003; Okey *et al.*, 1989; Pohjanvirta *et al.*, 1998; Poland *et al.*, 1994), in particular the reported impaired signaling by the human AHR in a non-human cellular background attributed to co-factor incompatibility issues (Moriguchi, *et al.*, 2003). Most importantly, we tested the complete Neanderthal AHR mRNA sequence in a human instead of a rat cellular background. Furthermore, the ancient sequence tested included the 3’- and 5’-untranslated region (UTR) comprising 4 instead of 2 single nucleotide variants while leaving the original codon usage intact. Thus we obtained a very different result: no physiologically relevant difference in CYP1A1 transcription activation between the ancient and modern AHR variants, which disagrees with a major role for the AHR in the evolution of smoke detoxification.

## MATERIALS AND METHODS

### Detailed ancient AHR variant testing strategy

Hubbard *et al.* (2016) tested the protein-coding region of the AHR mRNA applying mammalian codon optimization. In addition, UTR regions of a gene transcript may contain various functional elements embedded in the primary sequence, or hairpins or other secondary structures, serving for example as protein or microRNA binding sites that play a role in translation initiation and regulation, mRNA stability, transport from or to the cell nucleus, or cellular localization (Hinnebusch *et al.*, 2016; Mayr, 2017). Specifically for the AHR mRNA a 5’-UTR variant has been described in the modern human population (rs7796976) that increases AHR mRNA expression (Prager *et al.*, 2016) and was also observed to occur in the ancient hominin AHR sequences studied here (position 185 in Fig. 1). For the 3’-UTR of the AHR mRNA a binding site for microRNA-124 was recently described (Liu *et al.*, 2018) shown to be involved in regulating its expression level as well. Furthermore, changing the natural codons, especially for amino acids at positions where the compared AHR variants differ (Ala, Val, Arg, Lys), may affect the protein translation efficiency and the ultimate expression level as compared to the natural mRNA sequence (Brule *et al.*, 2017). Therefore, we diverted from the strategy followed by (Hubbard *et al.*, 2016), in that we tested the complete and original AHR mRNA sequence, including the 5’- and 3’-UTR without applying mammalian codon optimization.

**Fig. 1.**
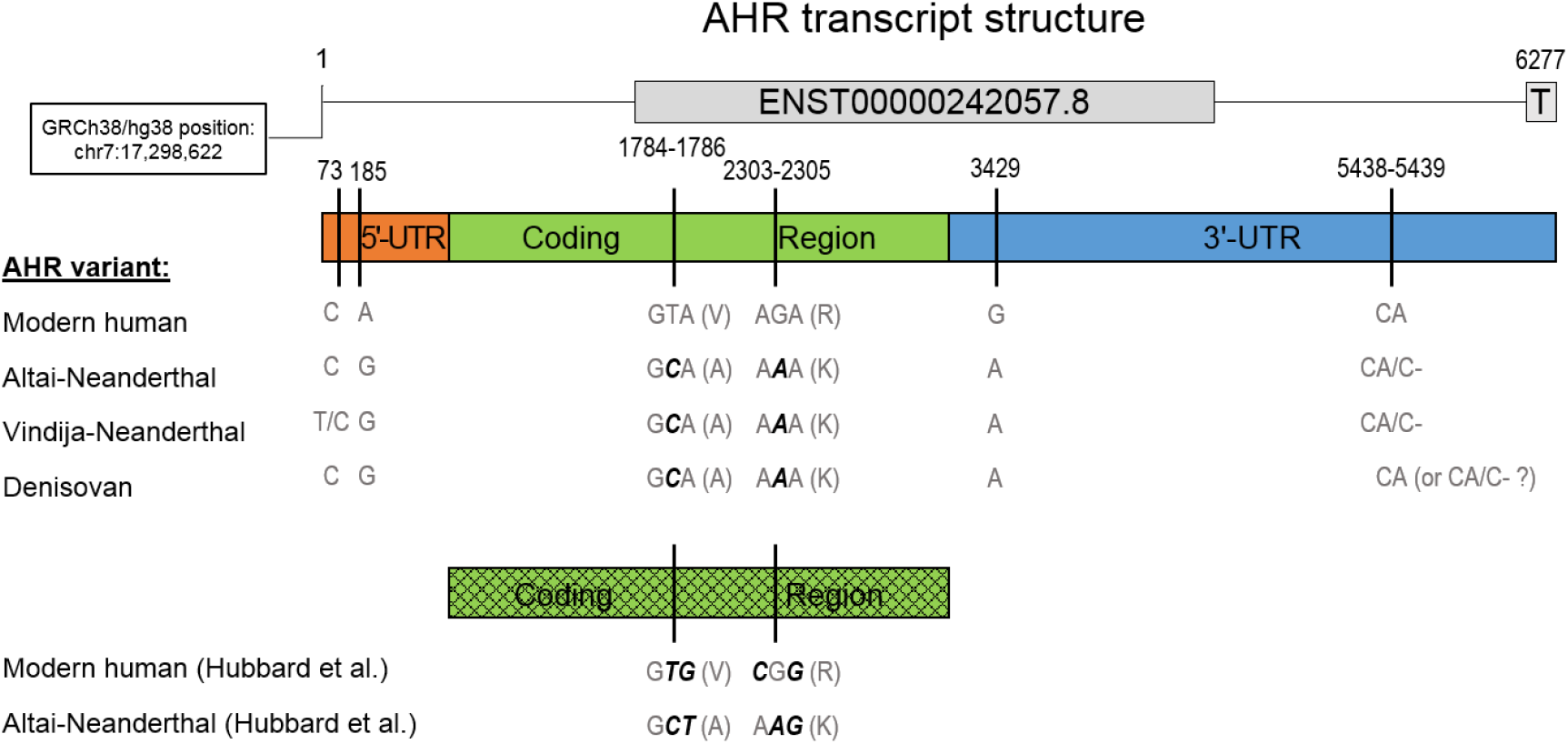
Map of the modern human and ancient hominin AHR mRNA sequence variants studied here and by Hubbard et al. (2016). The modern human AHR variant tested was based on manually expert-curated sequence information from the Ensembl and RefSeq databases (Materials and Methods). The bar gives a scaled representation of this 6277 bp sequence including the 5’-untranslated region (UTR), the AHR protein-coding region, and the 3’-UTR. The numbers above are the coordinates of the variant sites within the Ensembl AHR reference transcript ENST00000242057.8. Below it the base, or codon plus encoded amino acid (V = valine; A = alanine; R = arginine; K = lysine) present in each AHR variant is indicated. Cross-hatching indicates optimization for mammalian codon use and minimal secondary mRNA structure; absence of hatching indicates that the codon usage from the human reference sequences is maintained. Codon bases in bold italics differ naturally, respectively (in the synthetic sequences tested by Hubbard et al. 2016), have been changed as compared to the modern human reference. Apart from the indicated variable sites the reported ancient AHR transcript sequences are identical to the modern reference sequence.

### Synthesis of modern human and Neanderthal AHR cDNA sequences

The modern human AHR variant tested (Fig. 1) was the Ensembl AHR transcript AHR-201 (ENST00000242057.8) which is based on expert-curated sequence information and has been assigned the “gold” status, implying that this transcript is identical between Ensembl automated annotation and VEGA/Havana manual curation [http://jul2018.archive.ensembl.org/Help/View?id=151; <u>accessed Oct. 16, 2018].</u> An extra T deoxynucleotide was added to its 3’-terminus as reported by the RefSeq AHR mRNA record NM_001621.4, which is also manually curated [https://www.ncbi.nlm.nih.gov/refseq/about/; accessed Oct. 16, 2018].

The Altai- and Vindija-Neanderthal AHR mRNA sequence were retrieved from the chromosome 7 VCF data files published (Prüfer *et al.*, 2017; Prüfer *et al.*, 2014), and the differences with the modern AHR transcript AHR-201 found at six locations were determined as described in the Supplementary Data (SD), section SD1. The Altai-Neanderthal sequence carrying four single nucleotide changes as compared to the modern human reference was selected to be tested, as it turned out to be most representative for the Neanderthal and Denisovan AHR sequences known to date. A synthetic double-stranded copy DNA (cDNA) of the modern human AHR mRNA reference sequence (SD2) and of the Altai-Neanderthal derivative generated by site-directed mutagenesis (SD3), were purchased from BaseClear (Leiden, The Netherlands). Both synthetic sequences were verified to be 100% correct by Sanger sequencing.

### Construction of AHR expression constructs

The synthetic modern human and Altai-Neanderthal cDNA sequences were produced including 5’-terminal PstI and 3’-terminal XhoI restriction sites which were used to insert them into the expression vector pcDNA3.1/Zeo(+) (Invitrogen, Fisher Scientific, Landsmeer, The Netherlands) using standard recombinant DNA techniques (Sambrook *et al.*, 2001). These expression constructs were propagated in *Escherichia coli* DH5α and plasmid DNA was isolated using the E.Z.N.A. Endo-Free Plasmid DNA Mini Kit II (Omega bio-tek, VWR, Breda, The Netherlands) to obtain endotoxin-free plasmid DNA. For unknown reasons, specifically for the Neanderthal variant plasmid, the best yield was obtained when starting from a single, freshly grown bacterial colony and doubling the normal ampicillin level during propagation to 200 μg/ml. The identity of the obtained plasmid preparations was confirmed again by Sanger sequencing around the Ala381Val variation site (Fig. 1, C/T at position 1785) using the primers listed in SD4.

### Human cell culture and transfection

The human cervix epithelial adenocarcinoma cell line HeLa was purchased from the American Type Culture Collection, and the liver carcinoma cell line HepG2 from the European Collection of Authenticated Cell Cultures (Sigma-Aldrich Chemie N.V., Zwijndrecht, The Netherlands). General cell culture supplies were all purchased from Gibco (Life Technologies Europe B.V., Bleiswijk, The Netherlands) if not indicated otherwise: HeLa cells were cultured in DMEM + 4.5 g/l D-glucose + L-glutamine + 25 mm HEPES without pyruvate (42430-025) to which MEM non-essential amino acids (11140-035) were added and 10% fetal bovine serum (Sigma, F7524, Sigma-Aldrich Chemie N.V., Zwijndrecht, The Netherlands). HepG2 cells were cultured in EMEM + Earle’s salts without L-Glutamine (21090-022) to which MEM non-essential amino acids (11140-035) were added and 10% fetal bovine serum (American Type Culture Collection, ATCC 30-2020, LGC Standards GmbH, Wesel, Germany).

For transfections HeLa and HepG2 cells were seeded at approximately 80% density in 96-well plates (Greiner Bio-One 655180). HeLa cells were transfected using the *Trans*IT-HeLaMONSTER Transfection Kit (MIR 2904) and HepG2 cells with *Trans*IT-LT1 Transfection Reagent (MIR 2304) applying the standard protocols (both from Mirus Bio LLC, Sopachem b.v., Ochten, The Netherlands).

### Exposure of transfected HeLa cells

Twenty-four hours after transfection, 3 replicate wells with HeLa cells were exposed to each of the TCDD concentrations to be tested by adding an equal volume of medium containing twice the final concentration. As tetrachlorodibenzofuran (TCDF) used by Hubbard et al. (2016), TCDD is also a full AHR agonist with a 10 times greater potency to activate the AHR (WHO-TEF of TCDF is 0.1 (Van den Berg *et al.*, 2006)).

### AHR 5’-UTR - EGFP fusion reporter gene constructs

The modern human reference and ancient AHR mRNA sequence differ consistently at one position within the 5’-untranslated region (UTR), at base pair (bp) 185 according to the numbering of the human reference ENST00000242057.8. Both 5’-UTR variants were generated by PCR using Phusion High-Fidelity DNA polymerase (New England Biolabs M0530) and as a template the full-length AHR cDNA clone SC119159 purchased from Origene (delivered in vector pCMV6-XL4; distributor Acris Antibodies GmbH, Herford, Germany). A segment from this construct (bp 918-1612) carrying the complete SC119159 5’-UTR (185A-variant) and an upstream EcoRI restriction site was amplified by standard PCR procedures using the primer pair pCMV6-XL4_918-937 and AHR_ENST643-630_XhoI, the latter including a 3’-terminal extension with an XhoI restriction site and a 6 bp poly(A) clamp enhancing its digestion (see SD4 for sequence details and annealing temperature applied). The ancient G-variant of this segment was generated by overlap extension PCR (Heckman *et al.*, 2007). The upper mutated overlap segment (bp 918-1169 of SC119159) was generated using the primer pair pCMV6-XL4_918-937 and AHR_ENST200-170_185G_rev, and the lower mutated overlap segment (bp 1142-1612) with primer pair AHR_ENST173-203_185G_fwd and AHR_ENST643-630_XhoI. Before fusion PCR both segments were column purified using the GeneJET PCR Purification Kit K0702 (Thermo Scientific, K0702, VWR, Breda, The Netherlands) following the standard protocol. The fusion reaction of these overlap segments contained 100 pg of the short (252 bp) and 200 pg of the long fragment (471 bp; overlapping 28 bp with the shorter segment), 10 μM of the primers pCMV6-XL4_918-937 and AHR_ENST643-630_XhoI, 10mM dNTPs, 1x Phusion HF Buffer and 1 unit of Phusion DNA polymerase in 50 μl reaction volume. After initial denaturation at 98 °C for 30 seconds (sec) this reaction mixture was subjected to 5 cycles of 10 sec. 98 °C, 3 minutes (min.) 55 °C, and 30 sec. 72 °C, followed by a finishing incubation at 72 °C for 5 min. The undetectable amount of fusion product formed was column-purified using the GeneJET PCR Purification Kit and reamplified using the outer primer pair pCMV6-XL4_918-937 and AHR_ENST643-630_XhoI. The generated fusion product (707 bp) was column-purified and digested with EcoRI and XhoI and again column-purified, and then ligated applying standard cloning techniques into EcoRI and XhoI digested and subsequently gel-purified pcDNA3-EGFP reporter vector (AddGene, Cambridge MA, USA). The resulting construct was propagated in *Escherichia coli* DH5α according to standard methods (Sambrook, et al., 2001) and purified for transfection of human HepG2 cells using the E.Z.N.A. Endo-free Plasmid DNA Mini Kit II (Omega Bio-tek D6950-01, VWR, Breda, the Netherlands). The A- or G-variant type and insert integrity of the final plasmid preparations was confirmed to be 100% correct by Sanger sequencing. An aligned overview of the vector template, primers, overlap segments, overlap extension product, and AHR 5’-UTR target sequence aligned to the human AHR mRNA reference sequence is available in SD5.

### Quantification of AHR 5’-UTR-EGFP fusion protein expression

The 185A or 185G AHR 5’-UTR-EGFP reporter fusion constructs were co-transfected (ratio 1:1) with pBV-Luc (AddGene, Cambridge MA, USA), carrying a firefly luciferase coding sequence under control of a minimal promoter, into HepG2 cells known to feature natural AHR expression and signaling (Zeiger *et al.*, 2001). EGFP expression was measured using a Tecan 96-well fluorescence plate reader (Tecan Infinite M200 PRO) using the standard settings for EGFP quantification. EGFP expression was normalized for transfection efficiency in each well based on the independent, minimal promoter-driven luciferase expression conferred by the cotransfected pBV-Luc plasmid, which was quantified using flash kinetics and a double injector automated plate reader to subsequently inject luciferase substrate reagent (Flash Mix according to (Boerboom *et al.*, 2006) and 0.2 M sodium hydroxide stop reagent.

### Measuring mRNA expression levels using quantitative PCR

Dose-response curves for the induction of CYP1A1 mRNA expression by TCDD exposure were determined after 40-44 hours of exposure. Harvesting of cells from transfected and TCDD-exposed wells and subsequent quantification of mRNA expression levels by qPCR (quantitative polymerase chain reaction) were carried out using the Ambion Power SYBR Green Cells-to-C_T_ Kit (Thermofisher Scientific 4402954). Alternatively, for small scale experiments, total RNA from HeLa and HepG2 cells was isolated using TRIzol reagent (Invitrogen, Fisher Scientific, Landsmeer, The Netherlands), transcribed into cDNA using the iScript cDNA Synthesis Kit, and qPCR was carried out using IQ SYBR Green Supermix (both from Bio-Rad Laboratories B.V., Veenendaal, The Netherlands). Melt-curve analysis was performed to assure a single qPCR product of the expected melting temperature. A Bio-Rad CFX Connect thermocycler was used throughout.

### Data processing and statistics

Dose-response data were analyzed using GraphPad Prism 7 software to determine the best fit to the data of the equation Y = Bottom + [X * (Top-Bottom) / (EC_50_ + X)], describing the relation between the response (Y = CYP1A1 mRNA level) and the agonist concentration (X = concentration TCDD) for a receptor with one ligand binding site such as the AHR. The output includes best estimates for the basal (Bottom) and maximal CYP1A1 induction level (Top), and for the agonist concentration inducing a half maximal response (EC_50_) as well as a 95% confidence interval for these curve fitting parameters. Differences between these parameters for the modern human and Neanderthal AHR were considered significant (p < 0.05) when the output value for one AHR receptor variant laid outside the 95% confidence interval of the value calculated for the other AHR variant.

## RESULTS

### Differences between the ancient and modern AHR mRNA sequences

The splicing donor and acceptor sites and the 10 adjacent bases upstream and downstream of the 20 exon-intron borders of the modern human AHR reference transcript were found to be represented with 100% fidelity in the Altai- and Vindija-Neanderthal and Denisovan AHR genomic sequence, and were even very conserved from deeper down the primate lineage, justifying to assume no differences in the exon structure of the ancient hominin AHR transcripts. The AHR mRNA nucleotide sequences thus deduced from the ancient genomes studied here differed at 6 positions (Fig. 1) from the human reference genome (assembly GRCh38/hg38). The single nucleotide variants observed in the three ancient hominin sequences (Altai- and Vindija-Neanderthal and Denisovan) were almost identical: except for the heterozygous position 73 of the Vindija 5’-UTR where the C-allele was still matching the ancient hominin consensus and therefore chosen to be tested; furthermore, at position 5439 of the 3’-UTR the Neanderthals seem to have acquired a lineage-specific (A) deletion variant, since it is found in both the Altai- and Vindija-Neanderthal genomes (in the heterozygous state), whereas it seems absent in the Denisovan, and is very rare in the present-day human population (rs770112913 in dbSNP build 151: allele frequency = 0.0006). Hence, the 5438-5439 CA allele seems most representative for the available ancient genomes and was therefore studied here. SD6 shows an alignment of all sequences studied here and by Hubbard *et al.* (2016).

### Synthesis of cDNA copies of modern and ancient AHR mRNA sequences

A DNA copy (cDNA) of the complete modern human AHR reference transcript was produced by gene synthesis; the Altai-Neanderthal variant chosen as the representative ancient AHR mRNA sequence was generated from the modern AHR gene synthesis product by site-directed mutagenesis (see SD2 and SD3, respectively).

### Validation of the HeLa cell line as an AHR variant test system

The human cervix epithelial adenocarcinoma cell line HeLa has been reported as a cell system with very low endogenous AHR expression validated for testing AHR gene variants (Koyano *et al.*, 2005). HeLa cells were transfected with an expression construct carrying a cDNA sequence from the complete Altai-Neanderthal AHR or the modern human AHR reference mRNA, as detailed in Methods, with essentially no difference in transfection efficiency (Fig. 2). We also observed an up to 130 times increase in the AHR mRNA level upon transfection with the AHR expression constructs as compared to empty vector (data not shown).

**Fig. 2.**
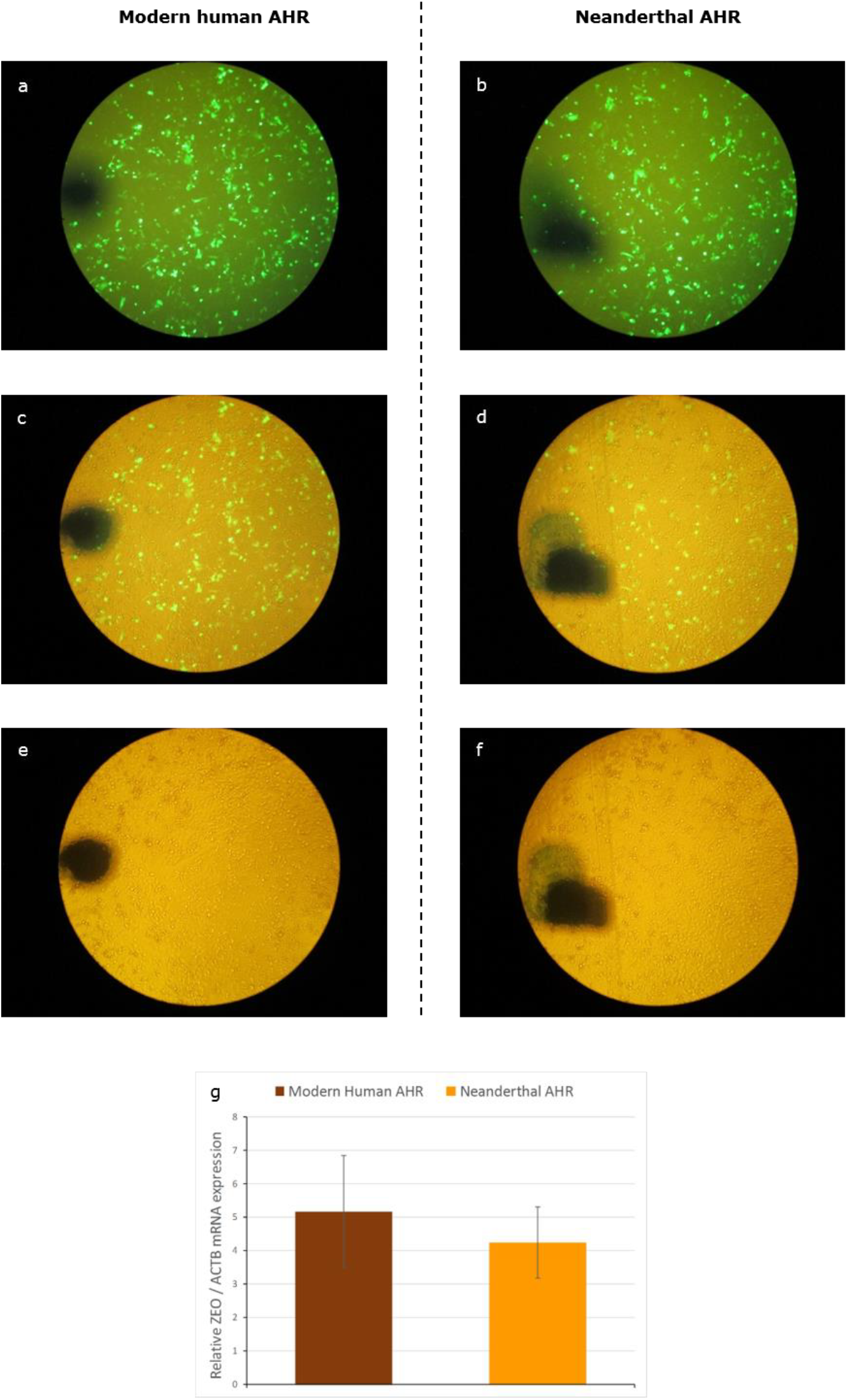
HeLa cells were transfected with a pcDNA3.1/Zeo(+)-based expression construct for the complete modern human (panel a,c,e) or the Neanderthal AHR mRNA (panel b,d,f). Panel a-f) Cotransfection (1:1 w/w) with the EGFP expression plasmid pcDNA3-EGFP to visualize the transfection efficiency; a,b) EGFP fluorescence; c,d) EGFP fluorescence + phase-contrast transillumination; e,f) transillumination only; the black marker spot is applied for orientation. The pcDNA3.1/Zeo(+) vector backbone is carrying a zeocin resistance gene expression cassette (ZEO). Therefore, ZEO mRNA expression is proportional to the number of functional construct molecules inside the cells upon transfection and allows to quantify the transfection efficiency. To correct for possible differences in cell numbers between transfections, the ZEO mRNA levels as quantified by qPCR were normalized for β-actin (ACTB) expression and the mean and SEM of the ZEO/ACTB mRNA expression ratio (n = 13) was plotted (panel g). A t-test on the difference in mean ratio between the transfections with the modern human AHR and Neanderthal AHR expression constructs showed no significant difference (p < 0.648).

Urged by preliminary observations that HeLa cells also showed CYP1A1 induction upon transfection with the AHR expression constructs without co-transfection with an Ah receptor nuclear translocator (ARNT) expression construct, we measured the endogenous ARNT mRNA level in HeLa and in HepG2 cells (Fig. 3), showing that the AHR-deficient HeLa cells expressed about 2 times higher endogenous ARNT mRNA levels as the notoriously AHR-competent human hepatocellular carcinoma cell line HepG2 (Zeiger, *et al.*, 2001), hence confirming that ARNT levels were not limiting AHR-mediated responses in these HeLa cells. Therefore we omitted the cotransfection with ARNT in our main experiments in contrast to the original protocol (Koyano, et al., 2005).

**Fig. 3.**
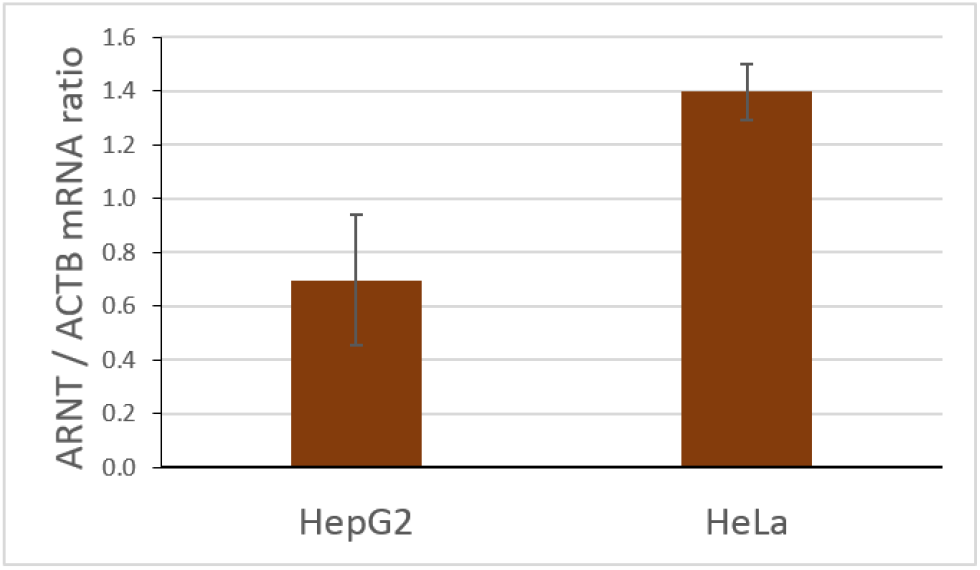
ARNT mRNA expression level in AHR-deficient HeLa as compared to AHR-competent HepG2 cells as measured by qPCR.

### CYP1A1 induction in AHR-neanderthalized human cells

The transfected HeLa cell population was exposed to a 0 – 10000 pM TCDD concentration range and the dose-response curve for induction of CYP1A1 mRNA expression was determined (Fig. 4).

**Fig. 4.**
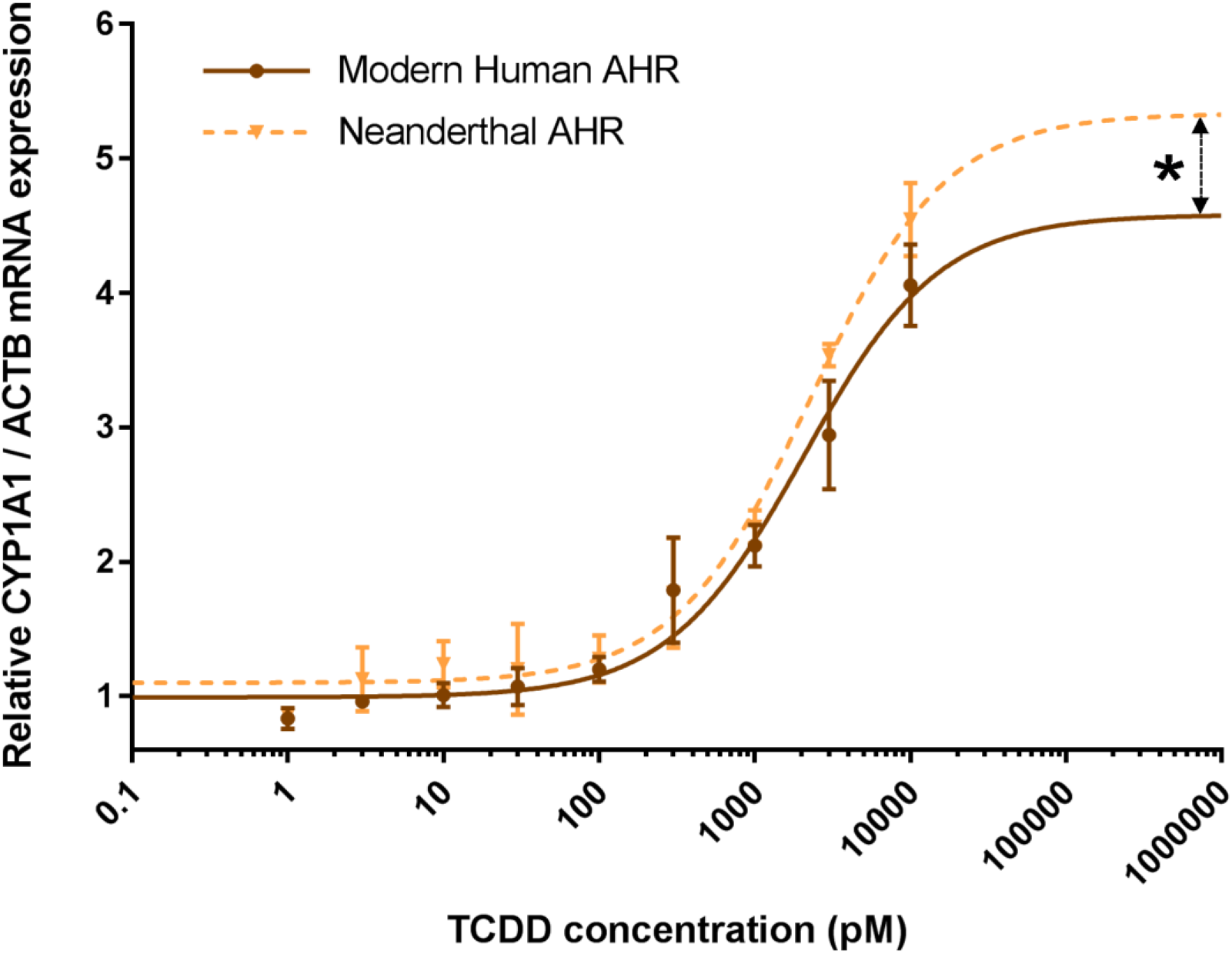
Dose-response relation observed for TCDD-induced CYP1A1 mRNA expression in HeLa human cervix epithelial adenocarcinoma cells transfected with an expression construct carrying a cDNA copy of the modern human or Neanderthal AHR mRNA, respectively. CYP1A1 and β-actin (ACTB) mRNA expression was measured using qPCR after 40-44 hours of exposure to TCDD. CYP1A1 expression was normalized for ACTB expression. The CYP1A1/ACTB ratios where then normalized to the basal level for the modern human AHR and all other values expressed as a fold value relative to this value set at 1, and plotted against the TCDD concentration in the culture medium during exposure. Effective concentration 50% (EC_50_), basal and maximal values for TCDD-induced CYP1A1 mRNA expression were calculated by fitting the one-site receptor-ligand binding equation (see Methods) to triplicate experimental data, which showed that only for the extrapolated maximal CYP1A1 expression level there was a statistical significant (p < 0.05) difference between the cells transfected with the modern human and the Neanderthal AHR, not for the basal level and EC_50_.

The curves for the cells transfected with the modern human and the Neanderthal AHR almost coincide. Upon fitting the one-site receptor-ligand binding equation (see Methods) to the data, only a small, but significant difference in the extrapolated maximal CYP1A1 induction was observed (Neanderthal AHR attains a 17% higher maximum), not in the basal level and the effective concentration 50% (EC_50_) (Table 1).

**Table 1.** Parameters of the dose-response curves for CYP1A1 induction by TCDD in HeLa cells transfected with the indicated AHR variant.

| AHR variant | EC<sub>50</sub><br>(nM) | Basal<br>CYP1A1/ACTB | Maximal<br>CYP1A1/ACTB |
| --- | --- | --- | --- |
| Modern human | 2.06 | 1.00 | 4.58 |
| Altai-Neanderthal | 2.28 | 1.10 | 5.34* |

\*) Significantly different ( $p < 0.05$ ) from HeLa cells transfected with a modern human AHR expression construct.

### Role of the 5’-UTR single nucleotide variation in the Neanderthal AHR mRNA

At position 185 the 5’-UTR of the modern human AHR carries a newly derived A variant (rs7796976 in dbSNP build 151: A allele frequency ≈ 0.21) which is unique for the human lineage among all presently sequenced primate genomes, whereas the Neanderthal and the Denisovan carry the ancestral G variant, and anatomically modern human (45 kya) from Ust’-Ishim (Fu, et al., 2014) was already heterozygous (see SD7).

To test the effect of both 5’-UTR variants on the expression of EGFP as a proxy for their effect on the protein expression directed by the respective AHR mRNA variants a 620 bp 5’-UTR sequence carrying each of these variants was attached to the enhanced green fluorescent protein (EGFP) coding region within an EGFP reporter gene construct (see Methods). These constructs were transfected into the human liver cell line HepG2, which is known to express all factors required for fully functional AHR signal transduction (Zeiger, et al., 2001), and hence will be capable to reflect the natural function of the AHR 5’-UTR. It was observed that the fluorescence signal tends to be approximately 1.3 times higher when the Neanderthal 185G 5’-UTR variant was inserted before the EGFP reporter (data not shown). This differential effect was not statistically significant but has been described before (Prager, et al., 2016), and is likely due to an effect on translation efficiency of the 185G variant, which could therefore have contributed to the significantly higher (1.17 x) maximal CYP1A1 induction observed under control of the Neanderthal AHR as compared to the modern human AHR.

## DISCUSSION

Since ancient hominin cells cannot be reconstructed to date, the “next best” and feasible option to obtain relevant functional information regarding ancient gene variants is expression in a modern human cellular background. The HeLa human cervical adenoma cell line is particularly suitable to carry out the comparison between the ancient and modern AHR, since it has been reported as valid for functional testing of AHR gene variants (Koyano, et al., 2005), because of a low background of endogenous AHR, and nevertheless expressing all essential components, such as ARNT (this study) to facilitate AHR signaling upon transfection with an AHR expression construct. Besides ensuring a homologous cellular background, our study compares the complete ancient hominin AHR transcript to the modern human reference, including the 5’- and 3’-UTR with 2 single nucleotide variations (SNVs) in addition to the 2 SNVs occurring in the coding region, while leaving the original codon usage intact.

Applying these test conditions, we observed that the Neanderthal AHR has a very similar trans-activating potency as compared to the modern human AHR: no significant difference in basal level and EC_50_ for induction of CYP1A1 mRNA, and only a small (17% higher) difference in the maximal induction level, found at very high TCDD concentrations, above 3 nM of TCDD toxic equivalents (TEQ). It is debatable, however, whether such high on-target concentrations are still relevant for real-life exposure levels. These results apply to the Denisovan as well because its AHR mRNA sequence is identical to the tested Altai-Neanderthal sequence (Fig. 1), and is likely to be representative for the Vindija-Neanderthal as well, since there is only a deviation from the Altai sequence at two UTR positions at the most, dependent on the allele considered and the unknown haplotype coupling of these differences, which have not been reported in modern humans (dbSNP, build 151) to be of any physiological relevance.

### Impact of the 5’-UTR variant

The higher maximal CYP1A1 target gene activation observed with the Neanderthal AHR might be due to the A > G substitution at position 185 in the 5’-UTR of the AHR mRNA which is likely affecting the translation efficiency of the AHR mRNA. The Neanderthal 185G 5’-UTR variant is still abundant in the present-day human population at a 79% frequency (dbSNP build 151, rs7796976), and was found associated with higher AHR mRNA expression and an epithelial barrier defect in smokers with Crohn’s disease (Prager, et al., 2016), which is consistent with the AHR expression trend that we describe here and shows that increased rather than decreased susceptibility to the adverse effects of smoke exposure is associated with the Neanderthal variant.

### Other genes playing a role in deactivation of environmental toxins

The new results presented here imply that the Ala381Val mutation that became fixed in the modern human lineage might be less relevant for AHR receptor activation than suggested by Hubbard et al. (2016) and not likely to have a major impact on the toxicity of dioxin-like compounds. Instead, many more genes have been shown to be involved in detoxification of dioxin-like toxicants (Moorthy *et al.*, 2015), which appeared to be predominantly in the ancestral, protective state in ancient hominins, and some of which gave rise to newly derived, less protective variants in the modern human lineage (Aarts, et al., 2016).

### Impact of the Ala381Val adaptation in the modern human lineage

In a human cellular background no difference in EC_50_ for CYP1A1 induction was found, and thus no difference in affinity between Ala381 and Val381 AHR protein variants for a typical AHR agonist such as TCDD, and that may be, in fact, not so surprising. The initial comparative studies on the human AHR and the AHR from dioxin-sensitive and -resistant mouse and rat strains showed that the observed length variation of the AHR polypeptide has an equally important impact on its functionality as the Ala>Val mutation (Ema, et al., 1994; Pohjanvirta, et al., 1998). On the other hand, the Ala>Val variation has only been studied in a rodent cellular background (Hubbard, et al., 2016; Okey, et al., 1989; Poland *et al.*, 1990), never in human cells, in which its effect may appear even less prominent. We hypothesize that the Ala>Val mutation, which is located on the edge of the ligand-binding domain, and in the immediate vicinity of the transactivation domain (Ema, et al., 1994), might influence the interaction with co-factors through an impact on the transactivation domain, rather than have a direct effect on the ligand-binding domain. In a heterologous cellular background, its effect on co-factor binding is likely to be different, and the altered overall conformation when in complex with heterologous cofactors could ultimately also affect the ligand binding equilibrium (Billas *et al.*, 2013; de Vera *et al.*, 2017). Consistent with this hypothesis are reports that the mouse AHR protein, in addition to several synergistic stimulatory subdomains, contains an inhibitory subdomain in its C-terminal trans-activating domain, which interacts with heat-shock protein HSP90 and functions in a species-, cell type-, and promoter-specific way (Ma *et al.*, 1995; Whitelaw *et al.*, 1994). This mechanism might explain why Hubbard et al. find a large difference in ligand binding affinity in a rat cellular background, which appears not to occur in a human cellular background as we show here for TCDD: the EC_50_ for the ancient AHR is only a non-significant fraction (11%) higher than the modern AHR value.

### Evolutionary mechanism behind the fixation event

Since our results imply no difference between the AHR from Neanderthals and modern humans concerning the activation by TCDD, the question arises why the Val381 variant has reached fixation in the modern human lineage. It is possible that there is still a significant physiological difference between these AHR variants towards other agonists than dioxin-like compounds or AHR antagonists, such as bioactive plant food components (Denison, et al., 2011), or endogenous AHR ligands such as tryptophan photo-oxidation products that need to be balanced at optimal levels (Wincent *et al.*, 2012). In that case, the Val381 AHR variant may confer a fitness advantage related to such AHR-active compounds, which could have driven the fixation process. Two other possible mechanisms that do not involve positive genetic selection are conceivable: 1. fixation by random genetic drift, which could have been enhanced by the small size of early modern human populations (Meyer *et al.*, 2012; Prüfer, et al., 2017; Whitlock, 2000); 2. positive selection on a gene that is genetically tightly linked to the AHR locus. A candidate could be the genomic region reported to be ranking at position 21 in terms of selective sweep strength in the modern human lineage, which is within 1 Mb from the AHR locus (chromosome 7: 18339385-18546684 in human reference genome assembly hg38; see Table S37 in (Green *et al.*, 2010)). This region contains one gene, HDAC9 encoding histone deacetylase 9. Histone deacetylation gives a tag for epigenetic repression and plays an important role in transcriptional regulation, cell cycle progression and developmental events (Delcuve *et al.*, 2012; Méjat *et al.*, 2005). In view of the close vicinity of the HDAC9 gene it is possible that, early in the modern human lineage, the AHR Val381 variant was driven to fixation as a result of the lack of recombination between the newly arisen AHR Var381 variant and the selective sweep region containing HDAC9.

### Implications for Neanderthal demise

As far as the deep history of fire use is concerned, our previous study (Aarts et al. 2016) was inconclusive regarding the use of fire by Neanderthals and Denisovans, as the high prevalence of low-risk, but mostly ancestral detoxification gene variants in their genome did not allow conclusions regarding positive genetic selection. Modern humans, on the other hand appeared to be evolving towards decreased detoxification efficacy in spite of their stronger dependence on fire use. The results presented here do not change this assessment, and are strongly at odds with suggestions that a mutation in the AHR may have given modern humans an evolutionary advantage over Neanderthals in adapting to smoke exposure.

### Future Directions

With a view to future research into the physiological properties of ancient hominin gene variants it appears crucial to reconstruct their expression conditions as faithfully as possible, especially the cell type used, which should be preferentially of human origin. We showed here that there is no essential difference between the ancient and modern AHR regarding activation by the toxic AHR agonist TCDD occurring in smoke. It remains to be seen, however, whether this indifference extrapolates to all classes of AHR-active compounds. Such studies, in particular of dietary and endogenous AHR agonists will be crucial to understand the full biological impact of the observed ancient AHR variations.

## SUPPLEMENTARY DATA DESCRIPTION

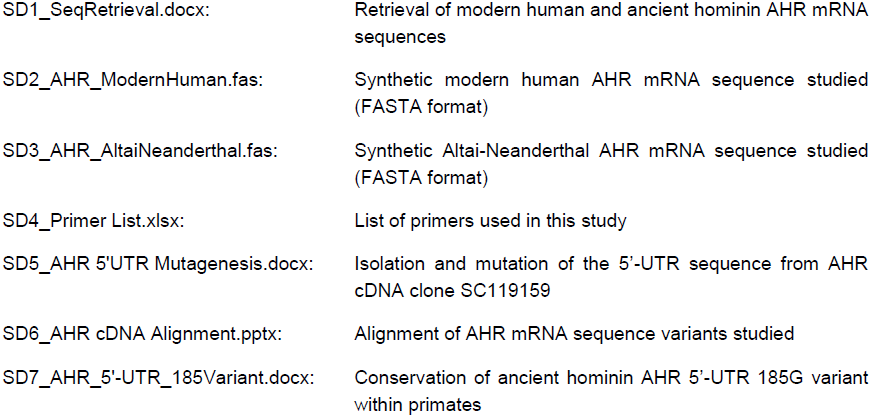

|  |  |
| --- | --- |
| SD1_SeqRetrieval.docx: | Retrieval of modern human and ancient hominin AHR mRNA sequences |
| SD2_AHR_ModernHuman.fas: | Synthetic modern human AHR mRNA sequence studied (FASTA format) |
| SD3_AHR_AltaiNeanderthal.fas: | Synthetic Altai-Neanderthal AHR mRNA sequence studied (FASTA format) |
| SD4_Primer List.xlsx: | List of primers used in this study |
| SD5_AHR 5'UTR Mutagenesis.docx: | Isolation and mutation of the 5'-UTR sequence from AHR cDNA clone SC119159 |
| SD6_AHR cDNA Alignment.pptx: | Alignment of AHR mRNA sequence variants studied |
| SD7_AHR_5'-UTR_185Variant.docx: | Conservation of ancient hominin AHR 5'-UTR 185G variant within primates |

## AUTHOR CONTRIBUTIONS

All four authors contributed to study design, interpretation of the results, and writing of the manuscript. In addition: J.M.M.J.G.A. conceived the study and carried out the experiments at the Laboratory of Molecular Biology of Wageningen University; G.M.A., a toxicologist retired from Wageningen University, in particular contributed with his toxicological expertise; H.J.F. contributed molecular-biological expertise, especially to design the AHR 5’-UTR-fused EGFP reporter gene study; W.R. is a Palaeolithic archaeologist responsible for the palaeoanthropological aspects of this study.

## ACKNOWLEDGEMENTS

The authors want to thank Mr. Jan W.G. Verver and Mr. Jan H.J. Hontelez from the Laboratory of Molecular Biology of Wageningen University for their excellent technical advice.

